# Large-scale computational modelling of the M1 and M2 synovial macrophages in Rheumatoid Arthritis

**DOI:** 10.1101/2023.09.11.556838

**Authors:** Naouel Zerrouk, Rachel Alcraft, Benjamin A. Hall, Franck Augé, Anna Niarakis

**Affiliations:** GenHotel—Laboratoire Européen de Recherche Pour La Polyarthrite Rhumatoïde, University Paris-Saclay, University Evry, Evry, France; Sanofi R&D Data and Data Science, Artificial Intelligence & Deep Analytics, Omics Data Science, 1, Av Pierre Brossolette 91385, Chilly-Mazarin, France; Advanced Research Computing Centre, University College London, London, UK; Department of Medical Physics and Biomedical Engineering, University College London, London, UK; Lifeware Group, Inria Saclay, Palaiseau, France

## Abstract

Macrophages play an essential role in rheumatoid arthritis (RA). Depending on their phenotype (M1 or M2), they can play a role in the initiation or resolution of inflammation. The M1/M2 ratio in RA is higher than in healthy controls. Despite this, no treatment targeting specifically macrophages is currently used in clinics. Thus, devising strategies to selectively deplete proinflammatory macrophages and promote anti-inflammatory macrophages could be a promising therapeutic approach in RA.

State-of-the-art molecular interaction maps of M1 and M2 macrophages in rheumatoid arthritis are available and represent a dense source of knowledge; however, these maps remain limited by their static nature. Discrete dynamic modelling can be employed to study the emergent behaviours of these systems. Nevertheless, handling such large-scale models is challenging. Due to their massive size, it is computationally demanding to identify biologically relevant states in a cell- and disease-specific context.

In this work, we developed an efficient computational framework that converts molecular interaction maps into Boolean models using the CaSQ tool. Next, we use a newly developed BMA tool version deployed to a high-performance computing cluster to identify the models’ steady states. The identified attractors are then validated using gene expression datasets and prior knowledge. We successfully applied our framework to generate and calibrate the first RA M1 and M2 macrophage Boolean models. Using single and double knockout simulations, we identified NFkB, JAK1/JAK2, and ERK1/Notch1 as potential targets that could selectively suppress proinflammatory macrophages, and GSK3B as a promising target that could promote anti-inflammatory macrophages in RA.

## Introduction

Rheumatoid arthritis (RA) is a complex inflammatory autoimmune disease whose aetiology is still not fully understood^1^. RA is primarily characterised by a persistent inflammatory cascade in the synovial tissue^2^, resulting in painful, swollen, rigid joints and, later, in extra-articular manifestations like gastrointestinal and cardiovascular diseases^3^. There is currently no cure for RA, and if prescribed treatments merely seek to reduce the inflammation and alleviate disease symptoms^4^, they have also been associated with various adverse events^5^. Recently, research has revealed that the innate immune system is crucial to initiating and developing RA pathogenesis^5^. Macrophages are one of the most common innate immune cell populations in RA, and their number significantly correlates with the disease severity^6^. Macrophage populations are heterogeneous and can differentiate into various phenotypes in response to the local microenvironment stimuli. The M1 and M2 phenotypes represent the extremes of their activation spectrum.

Consequently, depending on their phenotype, these cells play a role in both the initiation and resolution of inflammation^7^. The M1 macrophages are responsible for the overproduction of inflammatory cytokines and the release of matrix degradation enzymes, leading to cartilage destruction ^5^. They can also attract proinflammatory T cells and induce their hyperactivation. On the other hand, the M2 macrophages alleviate inflammation via 1) the production of anti-inflammatory cytokines, including IL-10 and TGF-β, 2) tissue homeostasis and repair ^8^, and activation of regulatory T cell functions ^9^. Due to their excessive activation and proliferation and enhanced anti-apoptosis ability, the proportion of M1 macrophages is higher than that of M2 macrophages in RA^6^. Two approaches currently exist for targeting macrophages: downregulating M1 phenotype and expanding M2 phenotype or repolarising M1 to M2 macrophages^10,11^. Despite this, no medicines specifically targeting macrophages are currently used in clinics^6,8^. Thus, understanding the specific approach for targeted depletion of the inflammatory macrophage while sparing other macrophage subsets and reestablishing macrophage balance might be a practical therapeutic approach in RA^12^.

Investigation of such complex diseases has been hindered by reductionist approaches, focusing on specific cellular components but failing to provide a global picture of the pathogenic mechanisms under study. Indeed, each molecule can rarely be assigned a distinct role^13^. Instead, cellular functions and phenotypes arise from the interactions between the biological system components^14^. Ongoing developments in high-throughput experimental techniques provide tremendous data regarding these molecular interactions. One strategy to represent them is their abstraction to networks ^13^. Initiatives have been conducted by creating mechanistic molecular interaction maps for various diseases^15–19^, including RA^20^. These maps are a rich source of knowledge. However, they remain limited in predictions and hypothesis testing due to their static nature.

One of the primary goals of dynamical modelling is to understand the emergent features and behaviours of such complex biological systems^14^. Several modelling approaches are available and can be divided into two categories: quantitative and qualitative modelling. The quantitative modelling approach better characterises a system but requires kinetic data as well as a high number of parameters. Because many of these characteristics are unknown and difficult to determine in most systems, these models are limited in size^21^. An alternative approach, the Boolean formalism, can overcome this limitation. It is an abstraction of the system where each biomolecule can have two values: zero for inactivity and one for activity. Changes in the biomolecules’ values are defined by logical rules using the Boolean operators “AND”, “OR”, and “NOT”. The regulation of this state variable is given in a parameter-free way, making Boolean modelling a viable option for large-scale systems with unknown kinetic parameters^22,23^. When simulated, Boolean models can reach a stable configuration called an attractor. Attractors represent the model’s long-term behaviour and have been connected to biological phenotypes, making their computation a key point in Boolean models’ analysis^23^.

Building and analysing Boolean models for large-scale complex biological systems remains challenging. When the logical rules are manually defined, the generated models are usually smaller and do not entirely cover the biological systems described in the molecular interaction maps. When the models are inferred automatically from maps, they are closer representations of the systems^24^, but these large-scale models are more complex, including especially a much higher number of inputs. Considering that the size of the state space of a Boolean model is exponentially dependent on its node number (2^n^ states for n nodes), computing all of their attractors is computationally demanding^25^ and identifying biologically coherent states is difficult, especially in a cell or disease-specific context.

This work presents an efficient computational framework to build, analyse and validate the behaviours of large-scale Boolean models with hundreds of nodes and a significant number of inputs. The framework uses publicly available molecular interaction maps to automatically infer their corresponding executable Boolean models via the CaSQ tool^24^. Our approach enables the analysis of the generated models in a synchronous scheme using a new version of the BMA tool^26^ deployed to a high-performance computing cluster set-up. The framework identifies all the existing attractors of the models using parallel computing and then tests their coherence against gene expression datasets and prior knowledge. It computes a similarity score that describes the ability of the model to reproduce what is known in the literature and observed in datasets. We successfully apply our framework to generate and validate the behaviour of the first RA M1 macrophage and RA M2 macrophage Boolean models using their corresponding maps within the RA-Atlas^20^. Although the heterogeneity of macrophages in RA has not been fully uncovered, these models aim to cover the phenotypic diversity of macrophages through a phenotype-specific representation of their secreted cytokines/chemokines, stimulatory molecules, receptors, and transcription factors^12^. We used these validated models to investigate potential mono- and bi-therapies that specifically downregulate proinflammatory macrophages and promote anti-inflammatory macrophages in RA synovium. We perform in silico simulations to evaluate new RA drug combinations and propose potential therapeutic repurposing.

## Results

We illustrate in this section how we built and validated the first large-scale Boolean models describing the RA M1 and M2 synovial macrophages using their maps that are available in the RA atlas^20^. We also demonstrate how we used the calibrated models to investigate potential therapeutic options that would specifically eliminate inflammatory synovial macrophages and boost anti-inflammatory macrophages in RA synovium. We perform in silico simulations to evaluate new RA drug combinations and propose potential therapeutic repurposing.

### 1. Generation of the Boolean models of M1 and M2 macrophages in RA

The updated RA M1 macrophage molecular interaction map includes 601 components interacting via 405 reactions. The updated version of the RA M2 macrophage map comprises 513 components and 323 reactions (figure 1). Converting these two maps into executable Boolean models with CaSQ^24^ produced a network of 309 nodes, 75 inputs, and 562 interactions for the M1 macrophage and a network of 254 nodes, 57 inputs, and 430 interactions for the M2 macrophage.

**Fig. 1.**
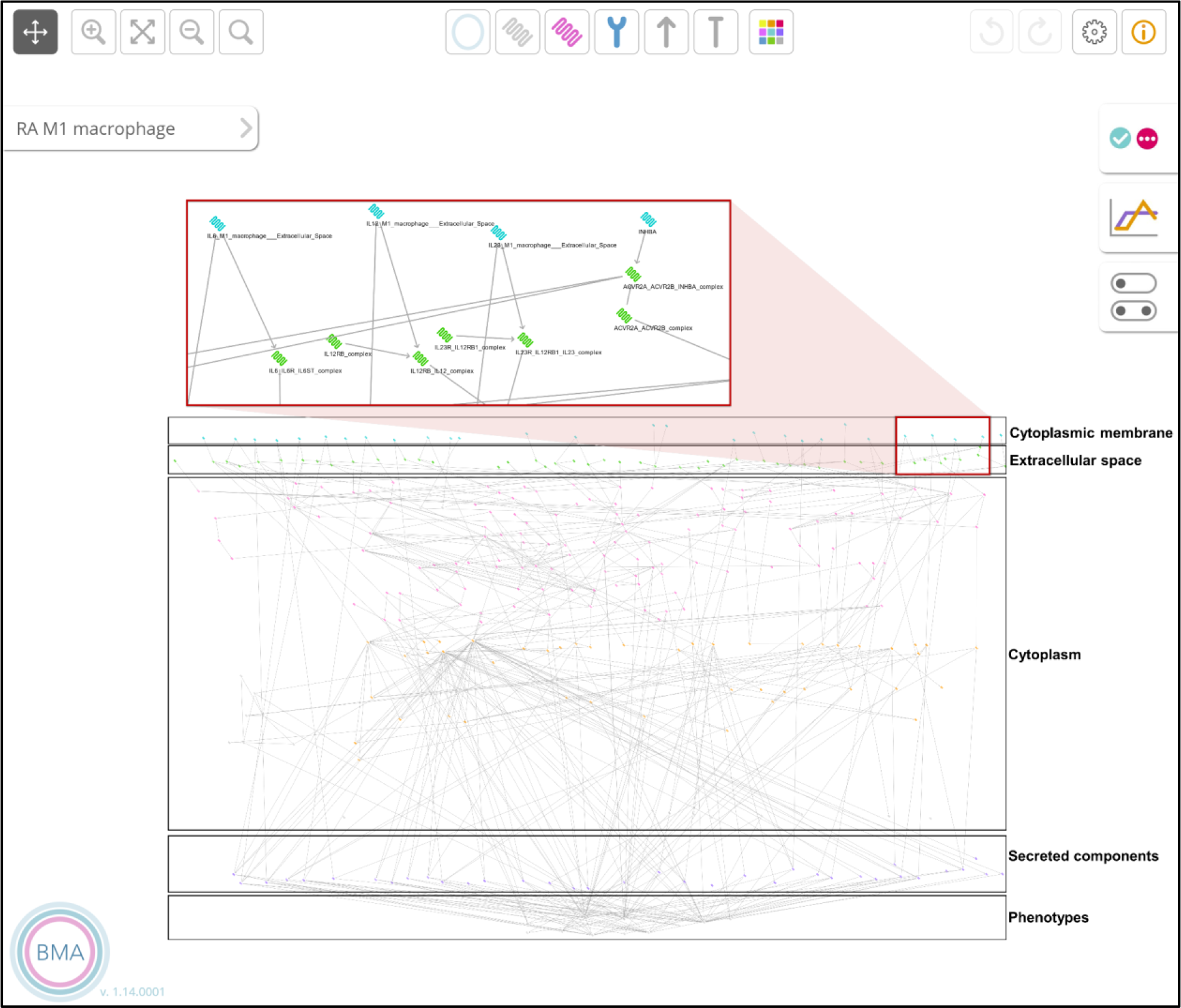
The RA M1 macrophage model in the BMA graphical interface

To focus on regulating the cell-specific phenotypes only, we used the export option in CaSQ via the argument -u to identify all the upstream nodes of these phenotypes.

Regarding the RA M1 macrophage model, the selected nodes are upstream of Apoptosis, Proliferation and Osteoclastogenesis phenotypes. The number of nodes decreased from 309 to 233, comprising 64 inputs. In the RA M2 macrophage model, the nodes of interest are upstream of Apoptosis and Proliferation phenotypes. Their number decreased from 254 to 169, including 39 inputs. The nodes that are not involved in regulating our phenotypes of interest were not considered. The inputs regulating these nodes were fixed at one, the default value in BMA.

### 2. Identification of differentially expressed molecules using literature search and transcriptomic data analysis

From both: the analysis of the GSE97779 dataset and an exhaustive literature search, we identified 105 and 88 differentially expressed biomolecules in the RA M1 and M2 macrophage models, respectively. The retrieved differential expressions can be associated with nodes at mRNA or protein levels in the models, assuming a linear relationship exists between the expression of mRNAs and the expression of their corresponding proteins. We discretised the differentially expressed molecules’ expressions: molecules that were overexpressed in RA were linked to the value 1, whereas molecules that were underexpressed in RA were linked to the value 0. Tables S-1 and S-3 list these differentially expressed molecules and their corresponding Boolean values in the M1 and M2 macrophages.

### 3. Computation of all the possible attractors of the models

Given the high number of inputs in the model, we reduced the list of inputs combinations by fixing the values of the differentially expressed ones. Based on the information displayed in Table S-1, 43 out of the 64 inputs present in the RA M1 macrophage model were fixed. The total number of input combinations was then equal to 2^21^. We used the BMA tool deployed to a machine with 96 single-core CPUs and 768 GB of RAM to run the attractors’ search. All the resulting attractors were steady states and were kept for further analysis.

Regarding the RA M2 macrophage model, 24 inputs were fixed using the information provided in Table S-3. The number of input combinations was then equal to 2^15^. All the corresponding attractors were steady states that we used for the following steps.

### 4. Validation of the models’ behaviours

First, we filtered the steady states according to the values of their cell-specific phenotypes. The biologically coherent Boolean values of these phenotypes were extracted from the literature in disease and cell-specific manner. They reflect the increased M1/M2 ratio in the synovial macrophage population and the enhanced osteoclastic bone resorption in the RA joint. Indeed, RA M1 macrophages predominate in RA synovial fluid due to their excessive proliferation (Proliferation phenotype in the model should be ON) and enhanced anti-apoptosis capabilities (Apoptosis phenotype in the model should be OFF) compared to the RA M2 macrophages (Apoptosis phenotype should be ON and Proliferation phenotype should be OFF in the model). In addition, the Osteoclastogenesis phenotype, which is only present in the RA M1 macrophage model, should be ON.

All the steady states of the RA M1 macrophage model passed through this filtering step, while only 8192 of the steady states of the RA M2 macrophage model did.

We calculated the similarity score between the list of differentially expressed molecules (Table S-1 and S-3) and their matching nodes in each filtered steady state. Regarding the RA M1 macrophage model, 384 steady states had the highest similarity score and their average vector was calculated. In the resulting vector, 222 nodes were fixed at zero or one, while eleven were not fixed (Table S-2). This model’s state can reproduce 99% of the observed Boolean values. Indeed, 104 of 105 differentially expressed nodes’ states matched their experimentally observed Boolean values. The only observed inconsistency is the CASP7 pro-apoptotic protein, which is overexpressed in RA macrophage samples but has a Boolean state equal to zero in the model.

Regarding the RA M2 macrophage model, 96 steady states had the highest similarity score. In their resulting mean vector, 158 out of 169 nodes were fixed at zero or one. Eleven nodes were not (Table S-4). This model’s state can reproduce 96,5% of the observed Boolean values. 85 out of the 88 differentially expressed nodes’ states matched their experimentally observed Boolean values. The only mismatches are BCL2L1 and MCL1, two upregulated anti-apoptotic proteins in RA macrophage samples with a Boolean state equal to zero in the model, and CASP3, a downregulated protein in RA macrophage samples with a Boolean state equal to one in the model.

### 5. Selective M1 macrophage depletion and M2 macrophage promotion in RA synovium

Selective downregulation of proinflammatory macrophages while promoting anti-inflammatory macrophages is a promising approach for inhibiting chronic inflammation and bone erosion in RA^10^. It can be achieved through the induction of the Apoptosis phenotype and the inhibition of the Proliferation phenotype in the RA M1 macrophage model and the activation of the Proliferation phenotype and the inhibition of the Apoptosis phenotype in the RA M2 macrophage model.

#### 5.1. Testing the effects of therapeutic targets’ single knockouts on the RA M1 and M1 macrophages

We performed an exhaustive search using the Therapeutic Target Database (TTD) to identify potential therapeutic targets (targets that have already been experimentally modulated) present in the models. It is a drug database designed to provide information about the known therapeutic protein and nucleic acid targets described in the literature, the targeted disease conditions, the pathway information, and the corresponding drugs/ligands directed at each target. The database currently contains 3578 targets and 38 760 drugs. Targets can be divided into four categories: successful targets, clinical trial targets, preclinical trial targets and research targets^27^. We screened the targets based on their associated drug’s Mode Of Action (MOA) and only kept the components that can be targeted by at least one inhibitor (1643 targets). Then, we identified the targets in the RA M1 and M2 macrophage models. Regarding the RA M1 macrophage model, 71 therapeutic targets that 988 drugs can inhibit were identified (Table S-5). Sixty targets that 1026 different drugs can suppress were identified in the RA M2 macrophage model (Table S-6).

We mimic the effect of these drugs using in silico knockout simulations. We use the calibrated state of both RA M1 and RA M2 macrophage models as initial simulation conditions (Table S-2 and Table S-4). Then, the models’ phenotype states after the target knockouts are compared to their corresponding calibrated states.

Table 1 summarises the identified therapeutic targets in both models. Inhibition of NFkB, in our models, stimulated the M1 macrophage’s death, inhibited the M1 macrophage’s growth (figure 2), and did not influence the M2 macrophage’s phenotypes. Even though ERK1 inhibition did not affect the M1 macrophage’s apoptosis, it did suppress their proliferation and reduce the release of most proinflammatory cytokines (CCL2, CSF2, IFNG, IL-18, IL-1, IL-6, and TNF). It also blocked the synthesis of the angiogenic factor VEGFA in the M2 macrophage model. GSK3B inhibition, on the other hand, induced the M2 macrophage’s proliferation while suppressing their apoptosis (figure 3).

**Table 1.** Single knockouts of the therapeutic targets from the TTD database that perturb the RA macrophages’ phenotypes.

| Target | Target type | Associated disease(s) | Drugs with the highest status | Effect on the RA macrophages' phenotypes |
| --- | --- | --- | --- | --- |
| NFkB | Successful | Irritable bowel syndrome, Rheumatoid arthritis, Choreiform disorder, Lupus erythematosus, Multiple sclerosis,... | Sulfasalazine (Approved) | -Induction of the M1 macrophage apoptosis<br>-Suppression of the M1 macrophage proliferation |
| ERK1 | Clinical trial target | Melanoma, Pancreatic cancer, Cancer, Arteries/arterioles disorder, Mature T-cell lymphoma. | BVD-523 (Phase 2) | Suppression of the M1 macrophage proliferation |
| GSK3B | Clinical trial target | Myotonic disorder, Acute myeloid leukaemia, Osteosarcoma, Fragile X chromosome, Myeloproliferative neoplasm,... | Tideglusib (Phase 2/3) | -Induction of the M2 macrophage proliferation<br>-Suppression of the M2 macrophage apoptosis |

**Fig. 2.**
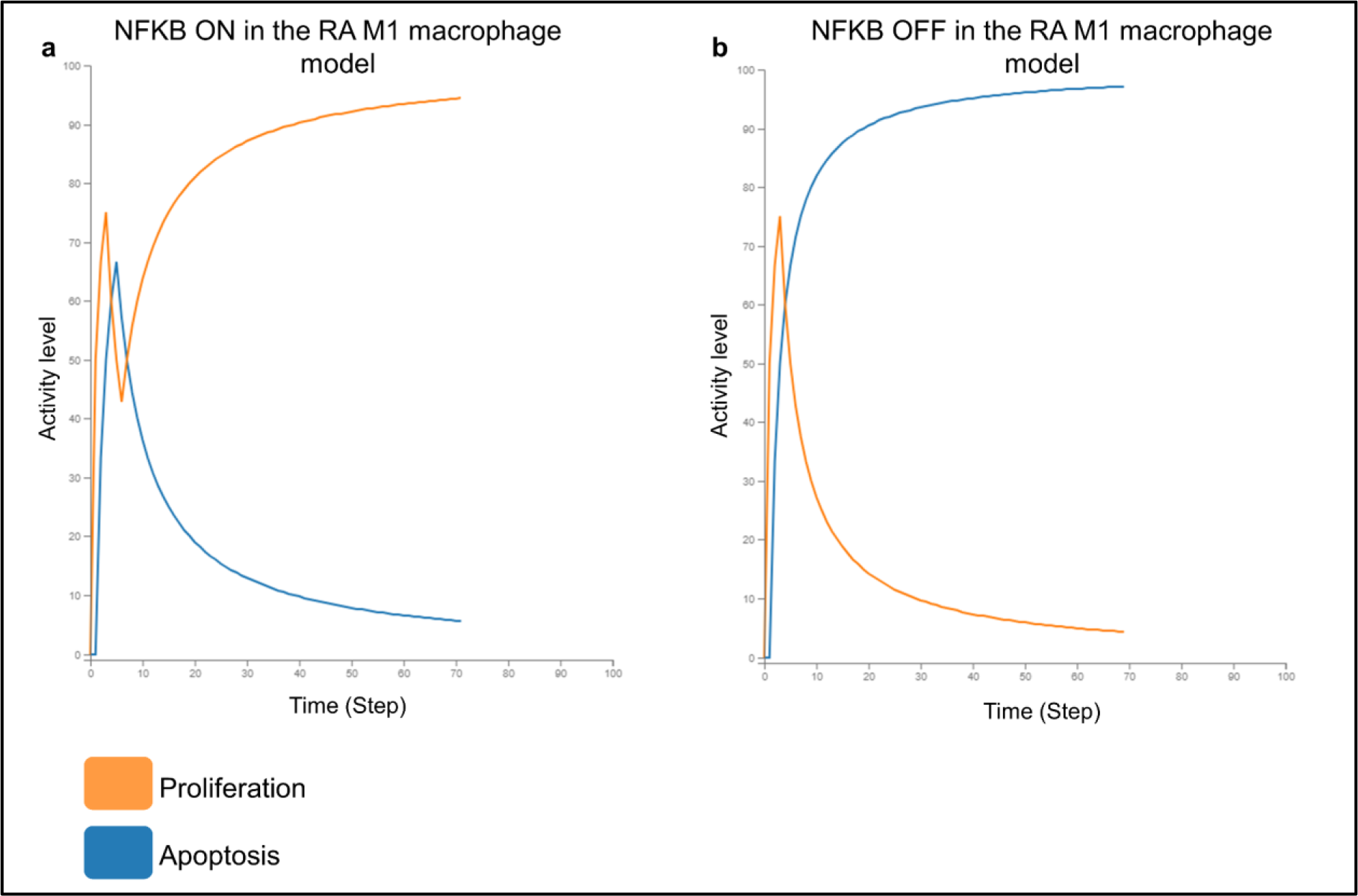
**a)** *In silico simulation of the proliferation and apoptosis phenotypes in the RA M1 macrophage model before* NFkB *KO. **b)** In silico simulation of the proliferation and apoptosis phenotypes in the RA M1 macrophage model after* NFkB *KO*.

**Fig. 3.**
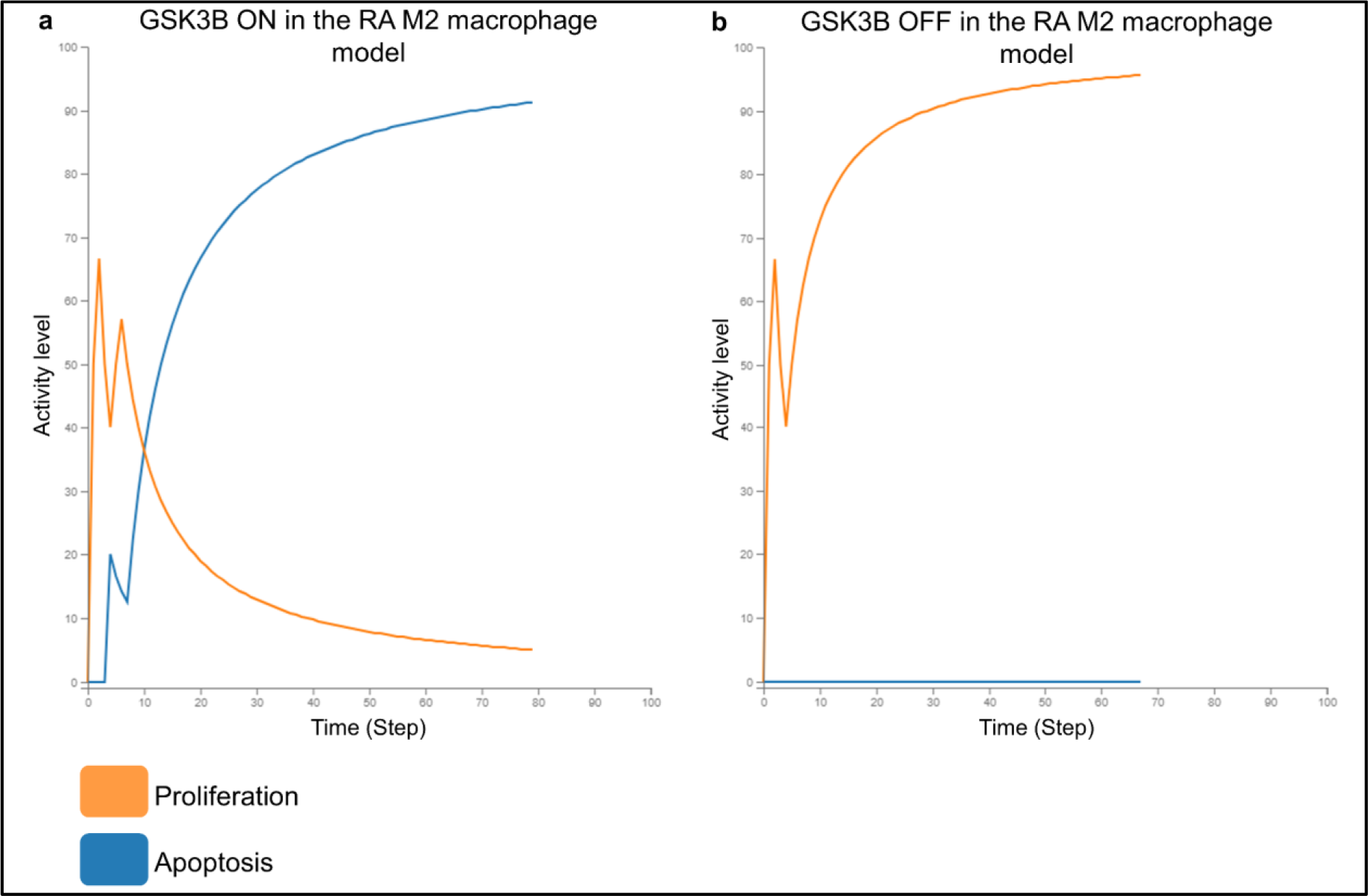
**a)** In silico simulation of the proliferation and apoptosis phenotypes in the RA M2 macrophage model before GSK3B KO. **b)** In silico simulation of the proliferation and apoptosis phenotypes in the RA M2 macrophage model after GSK3B KO.

#### 5.2. Testing the effects of therapeutic targets’ double knockouts on the RA M1 and M2 macrophages

In order to investigate the potential synergistic effect of the previously tested therapeutic targets on the models’ phenotypes, the targets were combined in pairs. Both RA M1 and RA M2 macrophages models were used to predict the outcome of their corresponding combined KO. We used the same initial conditions for the mono drug testing; then, we compared the perturbed states with their corresponding calibrated states.

Two thousand four hundred eighty-five drug combinations were tested using the RA M1 macrophage model. Among these combinations, the Notch1/ERK1 pair and the JAK1/JAK2 pair were identified as having a synergistic effect on the model’s phenotypes (table 2). Indeed, ERK1 KO alone inhibited the M1 macrophage’s proliferation. When combined with Notch1 KO, it also led to the promotion of the M1 macrophage’s apoptosis (figure 4). JAK1 and JAK2 separate inhibitions did not perturb the M1 macrophages’ phenotypes either. When paired together, they suppressed the M1 macrophages’ proliferation and induced their apoptosis. All the other drug pairs did not provide a synergistic effect on the model’s Apoptosis and Proliferation phenotypes. Apoptosis induction and proliferation suppression were only driven by NFkB KO.

**Table 2.**
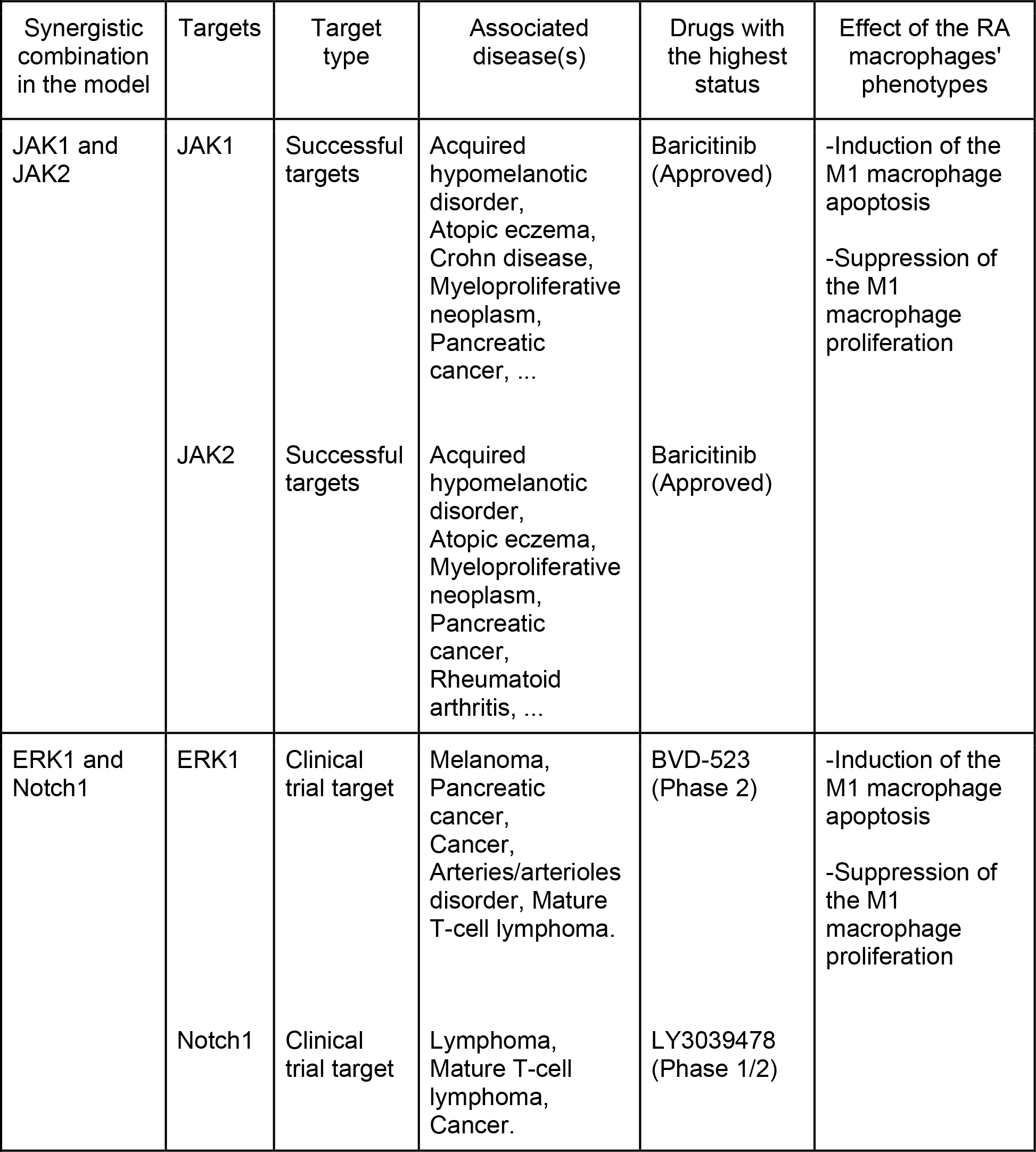
Combinations of therapeutic targets (from the TTD database) that perturb the RA macrophages’ phenotypes.

| Synergistic combination in the model | Targets | Target type | Associated disease(s) | Drugs with the highest status | Effect of the RA macrophages' phenotypes |
| --- | --- | --- | --- | --- | --- |
| JAK1 and JAK2 | JAK1 | Successful targets | Acquired hypomelanotic disorder, Atopic eczema, Crohn disease, Myeloproliferative neoplasm, Pancreatic cancer, ... | Baricitinib (Approved) | -Induction of the M1 macrophage apoptosis<br>-Suppression of the M1 macrophage proliferation |
|  | JAK2 | Successful targets | Acquired hypomelanotic disorder, Atopic eczema, Myeloproliferative neoplasm, Pancreatic cancer, Rheumatoid arthritis, ... | Baricitinib (Approved) |  |
| ERK1 and Notch1 | ERK1 | Clinical trial target | Melanoma, Pancreatic cancer, Cancer, Arteries/arterioles disorder, Mature T-cell lymphoma. | BVD-523 (Phase 2) | -Induction of the M1 macrophage apoptosis<br>-Suppression of the M1 macrophage proliferation |
|  | Notch1 | Clinical trial target | Lymphoma, Mature T-cell lymphoma, Cancer. | LY3039478 (Phase 1/2) |  |

**Fig. 4.**
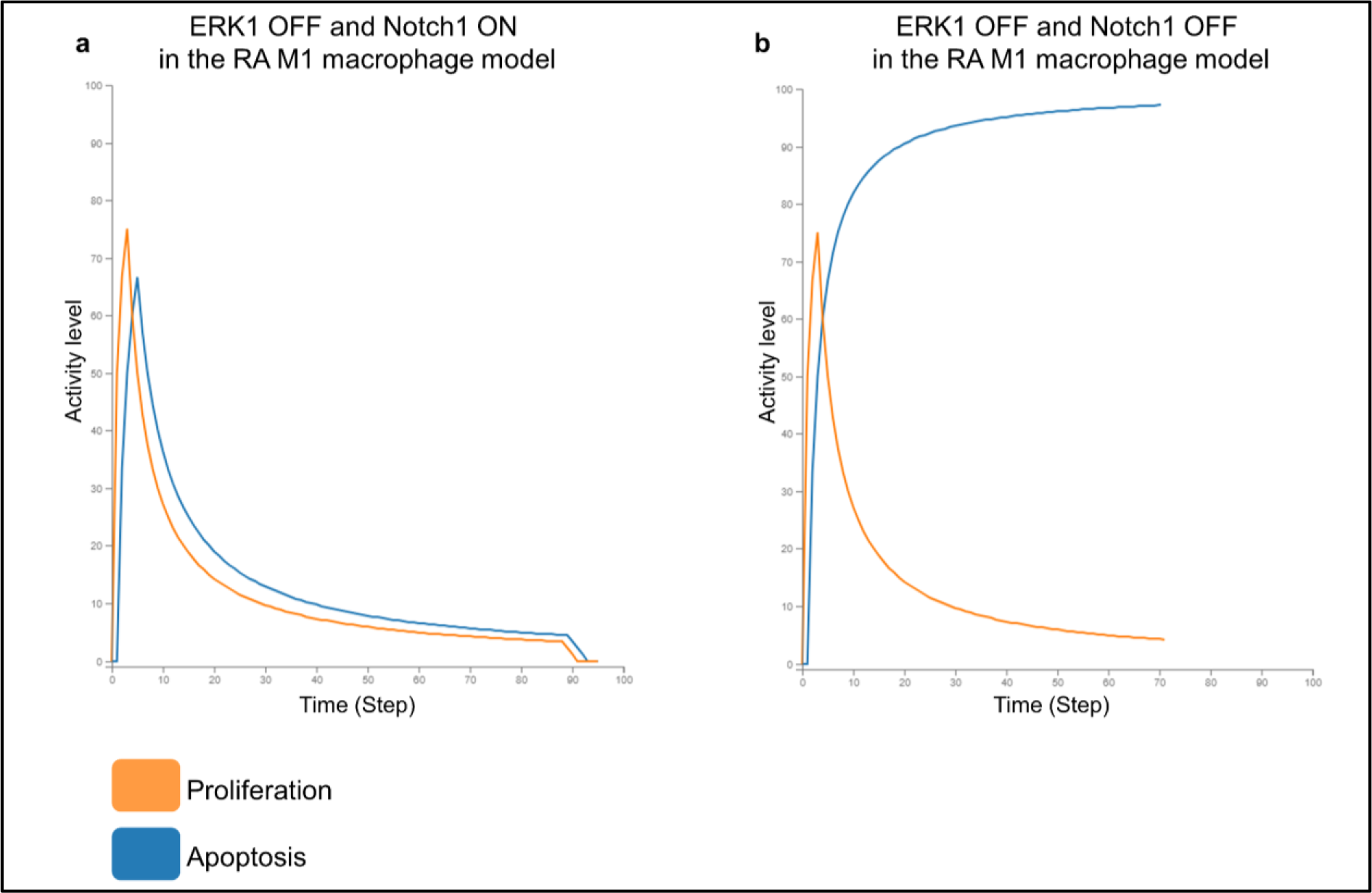
**a)** In silico simulation of the proliferation and apoptosis phenotypes in the RA M1 macrophage model after ERK1 KO. **b)** In silico simulation of the proliferation and apoptosis phenotypes in the RA M1 macrophage model after ERK1 and Notch1 KO.

Regarding the RA M2 macrophage model, 1770 drug combinations were tested. None of the drug combinations demonstrated a synergistic effect on the M2 macrophage model’s phenotypes. Apoptosis suppression and proliferation activation were only driven by GSK3B knockouts in the model.

#### 5.3. Testing of the effects of all possible receptor double knockouts in the RA M1 and M2 models

Receptors, located on both the cell surface and within the cell, are the molecular targets through which drugs produce their beneficial effects in various disease states. They are coupled to various signal transduction systems within the membrane and intracellularly and can therefore regulate responses to the cellular/tissue microenvironment^28^. In order to investigate the effects of double KOs of the cellular receptors present in the M1 and M2 macrophage models, receptors were combined two by two. Both RA M1 macrophage and RA M2 macrophage models were used to predict the outcome of their corresponding combined KO. We used the same initial conditions as for the mono and dual drug testing; then, we compared the perturbed states with their corresponding calibrated states.

Four hundred six double KOs were tested in the RA M1 macrophage model, while 300 double KOs were tested in the RA M2 macrophage model. None of these simulations perturbed the RA macrophages’ apoptosis or proliferation phenotypes.

## Discussion

The number of macrophages in inflamed synovial tissue overgrows during RA, and their polarisation plays a critical role in RA’s physiological and pathological progression^29^. Thus, selectively suppressing the M1 macrophages or boosting the M2 macrophage could be a promising strategy for treating RA. To investigate such complex mechanisms, we developed a framework to calibrate large-scale Boolean models that can be either automatically inferred from molecular interaction maps using the CaSQ tool^24^ or manually built in the BMA JSON format. We analyse the models using a newly developed BMA tool^26^ version that can be deployed on Linux-based high-performance computing clusters to identify all their steady states. Given the high number of inputs combinations in large-scale Boolean models, using HPC clusters enables high throughput model analysis. It overcomes the lack of computational power and takes advantage of parallel computing to considerably reduce the running time of the analysis.

We applied the proposed methodology to the large-scale RA M1 and M2 synovial macrophages interaction maps^20^, setting the path to many other disease maps to be explored, such as the Atlas of Cancer Signaling Network (ACSN)^17^, multiple sclerosis pathway map ^30^ or COVID19 disease map^19^. To analyse the resulting RA M1 and M2 macrophages models, we adapted the framework to make it relevant to the disease and cell type under study. Indeed, we filtered the models’ steady states according to the values of their cell-specific phenotypes and kept the ones that reflect the imbalanced M1/M2 ratio in the RA synovial macrophage population and the enhanced osteoclastic bone resorption in RA joints. In addition, to calibrate the models, we selected an RA and macrophage-specific gene expression dataset and carefully curated the extracted information from the literature to ensure that it was specific to both RA disease and synovial macrophages. The experimentally observed values we retrieved are cell and disease-specific but still contain a mix of M1 and M2 phenotypes’ expressions. Indeed, the identified mismatches in both models are related to biomolecules participating in the intrinsic and extrinsic apoptosis pathways. The M1 phenotype resists apoptosis, so pro-apoptosis components (CASP7) tend to be inhibited.

On the other hand, the M2 phenotype is pro-apoptotic; hence pro-apoptosis components are active (CASP3) while anti-apoptosis molecules are inhibited (BCL2L1, MCL1). The calibration of the RA macrophage models can also be performed against other datasets, either publicly available or proprietary. As RA is a highly heterogeneous disease, changes in certain DEGs are to be expected.

Because we are using Boolean formalism, the steady states of the models are binary vectors. To compare them to the differentially expressed molecules, we discretised the expressions. Biomolecules not differentially expressed, such as housekeeping genes, were not considered. Qualitative models with a higher granularity could be envisioned to address this limitation; nevertheless, additional computational resources would be needed to cope with the exponentially higher complexity of these models.

Most of the models’ nodes were set to zero or one in the selected steady states. The remaining nodes were not fixed due to a lack of information regarding their expression, meaning they may be found in both states (in different stable states). We could incorporate more layers of information by integrating other types of omics datasets, such as proteomic data, to set their Boolean values at either zero or one.

We performed in silico simulations on the calibrated models to investigate the effects of mono and bi-therapies on RA macrophage phenotypes. NFkB inhibition in our model led to selective suppression of the RA M1 macrophage. NFkB represents an interesting potential therapeutic target as it is a key transcription factor of M1 macrophages, responsible for the upregulated expression of M1 macrophage-derived cytokines in the RA synovium^31^. Several studies support the concept of NFkB inhibition for therapeutic interventions in inflammatory diseases ^32–35^. In RA, its in vitro inhibition induces apoptosis in fibroblasts, and contribute to a significant downregulation of M1 markers and upregulation of M2 markers^7,36^.

Further investigations showed that the observed beneficial effects of non-steroidal anti-inflammatory drugs (NSAIDs) and glucocorticoids, both used for RA treatment, are also due to NFkB inhibition^37–40^. However, their usage is limited due to severe side effects^40,41^ .Other NFkB inhibitors were identified, but most do not meet the standards to join clinical development programs^42–45^. Indeed, non-selective inhibition of NFkB in all cell types has multiple detrimental effects as it is critical for maintaining homeostatic cellular pathways. Biological treatments have been developed that directly target the products of NFkB-driven genes, such as TNF, IL-6 and IL-1. However, these treatments mainly target inflammation rather than apoptosis. Furthermore, as the mechanisms of apoptosis are highly sophisticated and several cytokines have synergistic biological activities^46^, inhibiting a single cytokine may not be optimal. This is further underlined by the simulations performed on our models that mimic such treatments (anti-TNF, anti-IL6, anti-IL1,…) and fail to induce apoptosis in the inflammatory RA macrophages. Therefore, the discovery of techniques for cell-type-specific NFkB inhibition is needed to shift the benefit/risk balance^47^.

Regarding the RA M2 macrophage model, GSK3B was identified as a promising target for promoting the M2 macrophage population in RA. GSK3B is involved in the progression of various diseases, including RA^48^. Evidence suggests that GSK3B plays a central role in signalling pathways relevant to macrophage function, including polarisation and inflammatory response^49^. Its inhibition in RA suppresses inflammatory responses in fibroblast-like synoviocytes and collagen-induced arthritis^50^. Furthermore, its inhibition in allergic rhinitis inflammatory disease increases the expression of the M2 phenotypic signature markers^51^. CREB1 is one of GSK3B’s targets ^52^. When GSK3B is inhibited, it induces CREB1 gain of function, sending an anti-inflammatory and anti-apoptotic survival signal in monocytes and macrophages^53^. It also increases M2 marker expression and promotes M2 phenotype in murine macrophages^54,55^.

We explored the synergistic effects that some therapeutic target pairs might have on macrophages models’ phenotypes. The M1 macrophage’s proliferation was suppressed by ERK1 knockout alone. When paired with Notch1 deletion, it also promoted the M1 macrophage’s death. The potential therapeutic value of co-targeting ERK1 and Notch1 has already been demonstrated in cancer but not RA. Indeed, it has been shown that targeting Notch1 enhances the efficacy of ERK1 inhibitors in cancer patients^56,57^. In RA, separate ERK1 and Notch1 inhibitions reduce inflammation in mouse collagen-induced arthritis^58,59^. Notch1 signalling, on the other hand, is known to regulate M1 macrophage fate through direct transcriptional and indirect metabolic regulation ^60^. We also identified the JAK1/JAK2 pair as a potential drug combination for the RA M1 macrophage depletion. Baricitinib, a Janus kinase (JAK) proteins inhibitor, is a Food and Drug Administration (FDA) approved for treating RA^61^. It prevents activation of STAT pathways and inhibits the cascade of transcription initiation of effector genes, which, in turn, prevents the autoimmune and inflammatory reactions associated with RA, including IFNg secretion. However, the way JAK inhibitors modulate macrophage phenotypes and whether this phenomenon explains their clinical benefit in RA is still not fully understood. A recent study showed that Baricitinib modulated the expression of membrane phenotype markers and the secretion of some cytokines in healthy macrophages^62^. Another study further supports the effect of JAKs inhibition on RA macrophage phenotypes by shifting the metabolic profile of M1 macrophage and rebalancing the metabolic reprogramming toward oxidative phosphorylation^63^.

Receptors are the molecular targets through which drugs produce their beneficial effects in various disease states. They are coupled to various signal transduction systems and can therefore regulate the cell’s responses to its microenvironment. We investigated the effects of double KOs of the macrophages’ receptors represented in the models, but none of the KOs perturbed the RA macrophages’ apoptosis or proliferation. These results underline the intricate crosstalks between the intracellular signalling pathways and the high synergistic activities of the various receptors represented in the M1 and M2 macrophage models.

Taken all together, these results further validate the behavior of our macrophage models through the identification of a new potential drug combination as well as targets whose potential/proved therapeutical benefit in RA has been highlighted in the literature.

The following steps of this work would be to combine the RA M1 and M2 macrophages maps with other cell-specific maps from the RA-Atlas^20^, namely the RA fibroblast and the RA Th1 maps, via the addition of intercellular interactions. Then, we would apply our framework to the resulting multicellular model. As the model’s size and complexity would considerably increase, we could assess our framework’s scalability.

## Methods

This section describes the computational framework we developed to convert publicly available molecular interaction maps into large-scale Boolean models, analyse their state space and validate their behaviours using prior knowledge and transcriptomic data (figure 5). This workflow is generalisable to manually built Boolean models in BMA JSON format. We also describe how we apply our framework to generate and validate the first RA M1 macrophage and RA M2 macrophage Boolean models using their maps that are available in the RA atlas^20^.

**Fig. 5.**
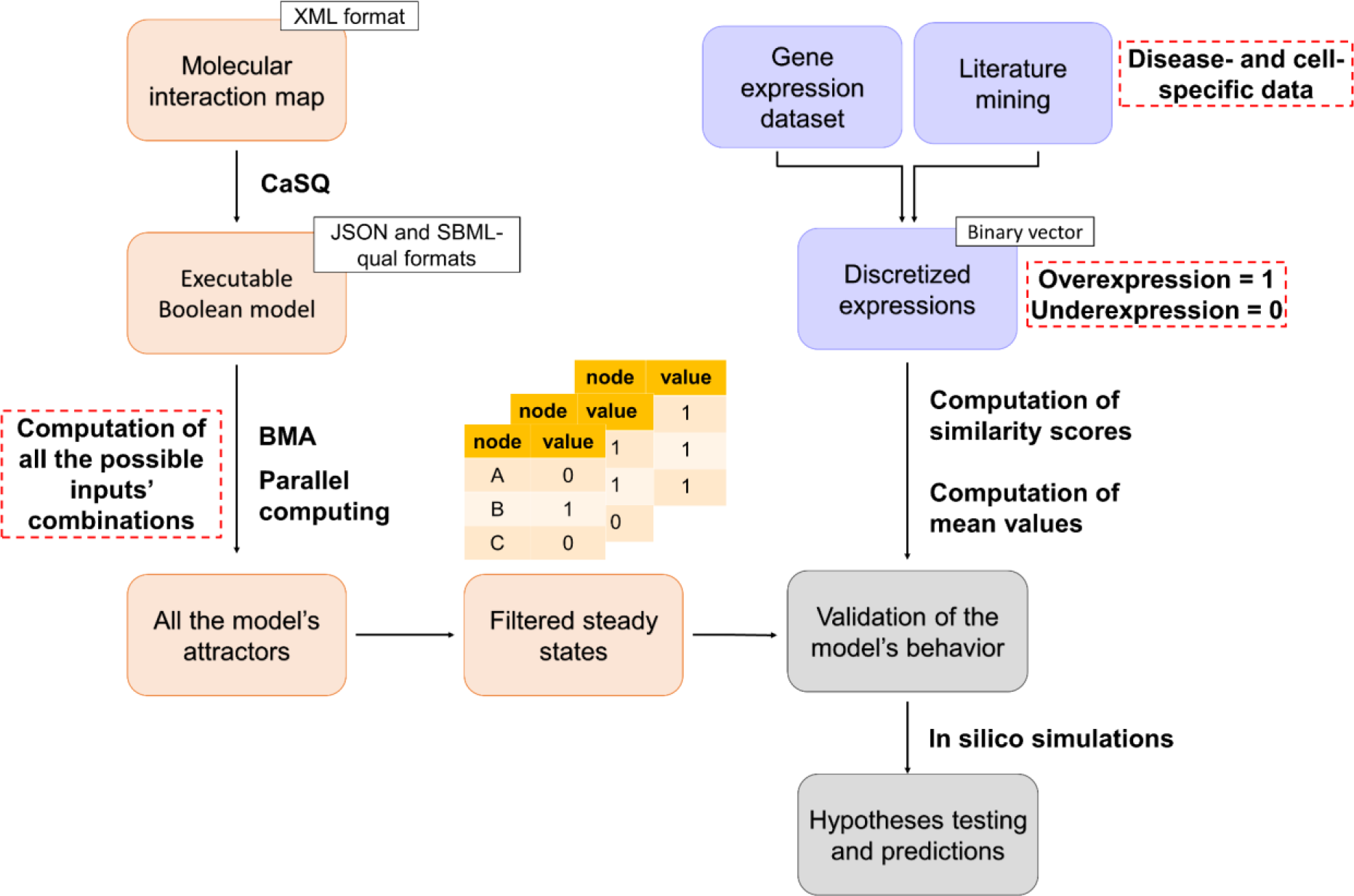
Schematic representation of the workflow we developed to generate and analyse large-scale Boolean models. Molecular interaction maps built in CellDesigner XML format are converted to executable Boolean models using the CaSQ tool. A new version of the BMA tool is then deployed on a high-performance computing cluster to identify all the models’ attractors. These attractors are filtered to keep only the steady states. Next, the filtered steady states are validated. Differentially expressed biomolecules in the models are identified using literature mining and transcriptomic data analysis. The identified biomolecule expressions are discretised and converted to a binary vector of experimentally observed Boolean values. After that, similarity scores are computed to describe the ability of the filtered steady states to reproduce the experimentally observed values. The steady states with the highest score are selected; their average vector represents the calibrated model’s state. The calibrated model can perform in silico simulations, test biological hypotheses and generate predictions.

### 1. Generation of Boolean models from molecular interaction maps

The workflow uses the CaSQ version 1.1.4^24^ to convert publicly available molecular interaction maps to executable Boolean models. CaSQ automatically infers logical rules based on the network topology and semantics for each node in the starting XML file. The tool produces either Systems Biology Marked up Language Qualitative (SBML-Qual)^64^ or BMA JSON executable files. Using the latter format, it is also possible to produce qualitative networks where nodes can vary over a wide range of discrete values, which is defined as granularity in BMA. Granularity defines the higher value the nodes can take in the model. Since we use Boolean formalism, the granularity used in the following analysis equals one.

We used their corresponding maps available in the RA-Atlas to generate the RA M1 macrophage and RA M2 macrophage Boolean models^20^. These two maps are built in the Systems Biology Markup Language (SBML) format^65^ using CellDesigner^66^ and are compliant with the Systems Biology Graphical Notation (SBGN)^67^. They cover cell-specific signalling pathways, gene regulations, molecular processes and phenotypes involved in RA’s pathogenesis. Biomolecules and reactions in these maps are manually curated and extensively annotated through PubMed IDs, DOI, GEO and KEGG identifiers, following MIRIAM (Minimum Information Required In The Annotation of Models) standards ^67^. Phenotypes are particular nodes in the maps. They describe biological states known to be active or inactive in RA. To make them more appropriate to the purpose of this work, we divided them into two categories. The first corresponds to cell-specific phenotypes, describing the cellular outcomes of RA synovial macrophages like proliferation and apoptosis. Depending on the map, their names end with “M1_macrophage” or “M2_macrophage” suffix. The second category is not specific to a particular cell type and corresponds to cellular signals and biological conditions in the RA joint, like inflammation and matrix degradation. Their names end with the ‘‘signal’’ suffix in both updated maps. We also looked for duplicates, removed them whenever found, and corrected the signalling pathways accordingly.

### 2. Stabilisation proof using Bio Model Analyzer (BMA)

BioModelAnalyzer (BMA) is a tool for constructing, analysing, and importing executable models of biological mechanisms^26^. The user is presented with a web-based interface, allowing for rapid and straightforward model construction and analysis. Whilst the GUI is the primary tool for interacting with BMA, a console tool is also available, giving access to a wide range of analysis algorithms and enabling scripting for large and complex combinatorial analyses. CaSQ can generate models in the BMA JSON format, which can be used with either version of the BMA tool.

Fundamental analysis in BMA is the proof of model stability that symbolically analyses the model attractors without explicitly calculating the model transitions. A modular proof algorithm is used under the synchronous update scheduler to show whether or not a single steady-state attractor exists and no cycles. Briefly, this proceeds in two steps. Initially, the ranges of individual variables are reduced to the set of reachable values by examining the input variable ranges and the target function. Stability is proven if this process reduces all ranges to a singleton and the global steady-state attractor is returned to the user. If this fails, it uses Boolean satisfiability (SAT) queries via a constraint solver, with the reduced variable ranges from the first step, to search first for multiple fix points (bifurcation) and then cycles. Finally, if neither cycles nor multiple fix points are found, the model must be stable and a final check searches for and returns the steady state^68^.

### 3. BMA architecture and underlying technologies

BMA is developed on the Microsoft .NET Framework and .NET standard, which tie the tools to Windows environments. The BMA web tool is hosted on Azure and is structured as two services, one hosting the user-facing client and another computing service dedicated to calculating proofs and simulations. The console tool is developed for Windows, which provides similar functionality to the compute service. To enable high-throughput model analysis and take advantage of parallelisation on high-performance computing facilities (typically Linux-based), we developed a prototype of the console tool based on the open source .NET core 3.1, which can be built using the dotnet SDK. All codes are available at 10.5281/zenodo.7541023.

### 4. Parallel computing for the calculation of all possible attractors

Attractors depend on the external stimuli the model receives from its environment. Stimuli in Boolean models are modelled in the form of inputs. Inputs are nodes with no upstream regulation. They are not associated with any logical rule in the model; therefore, their values are user-defined. In the BMA console tool, the user can assign values to the input nodes with the flag -ko that allows setting the specified nodes to be constants (zero or one). Depending on the input nodes’ state, the model reaches different attractors. To identify all the attractors of the model, we generate all the possible combinations of inputs’ values. For each inputs combination, we search for the corresponding attractor. We reduce the number of input combinations when possible by fixing the inputs associated with experimentally observed expressions. The Boolean values of these inputs are set based on the available literature and transcriptomic data (see Validation of the model’s behaviour section).

The computation time complexity is exponential. Indeed, the number of all possible combinations of inputs’ values equals 2^n^, n being the number of inputs that vary in the model. Given the high number of inputs in the inferred large-scale Boolean models, we failed to execute the attractors search on a local Windows machine with eight cores and 64 GB of RAM. Therefore, we deployed BMA on a high-performance computing cluster to compensate for the lack of computational power. The cluster capacity should be selected based on the model’s size and the number of input combinations to process. Figure 6 illustrates the number of input combinations the BMA console tool can process per hour using one core and various model sizes. As the attractor search is slower on larger models, the number of processed combinations decreases proportionally with the model size. Therefore, figure 6 can be used to estimate the computational resources required to execute the analysis depending on the model’s size. We utilised the Joblib python package as well ^69^ to parallelise the process and considerably reduce the running time of the framework.

**Fig. 6.**
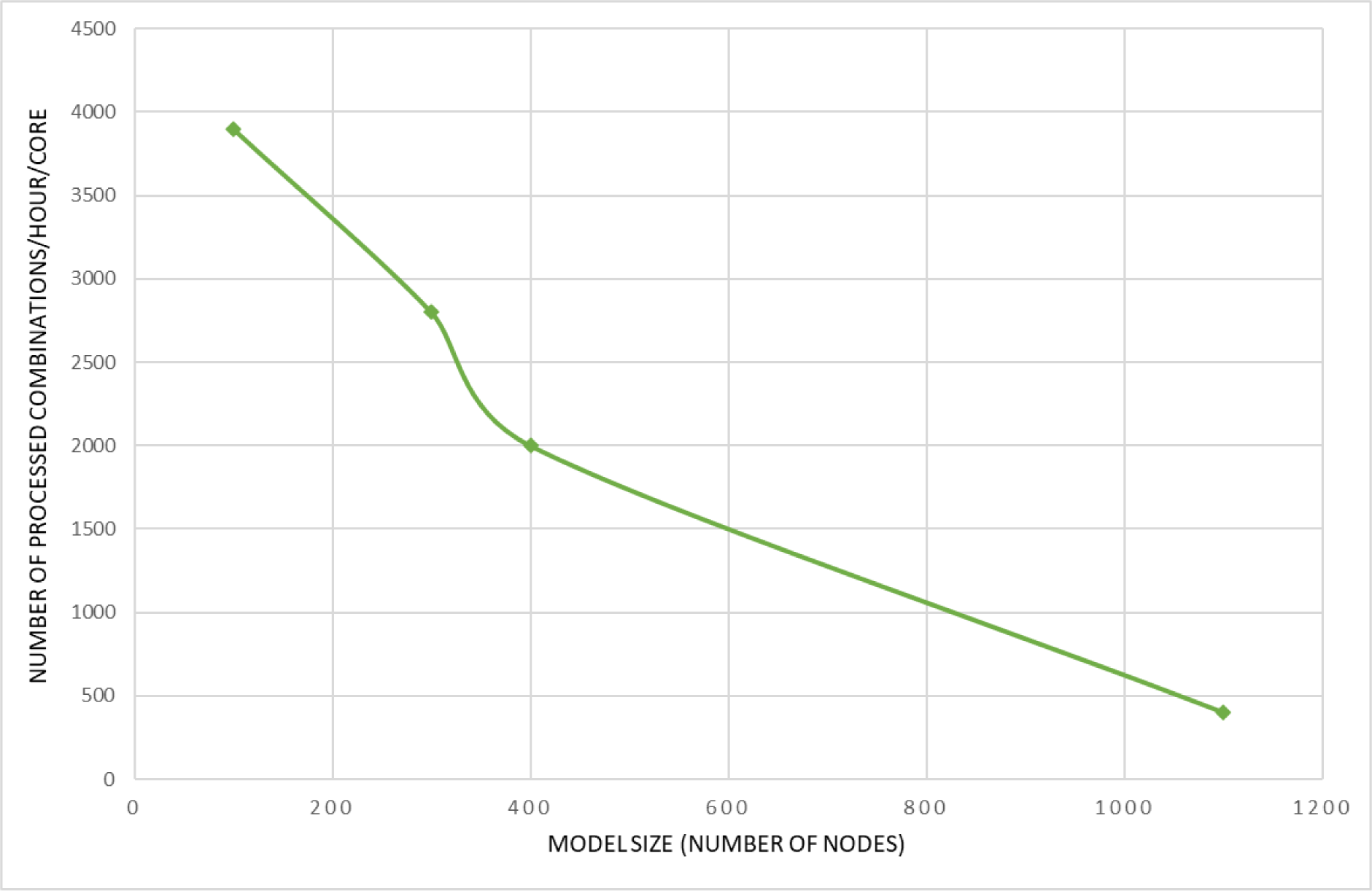
Plot showing the number of processed inputs’ combinations by BMA per hour using a single core machine.

### 5. Filtering the model’s attractors

While cycles are linked with oscillations such as the cell cycle, steady states are characterised by stable patterns of biomolecule activities; they can therefore be compared with gene expression datasets. For this reason, we filter the model’s attractors to keep only the steady states.

### 6. Validation of the model’s behaviour

The model’s state is validated based on the signal propagation from the inputs to the internal nodes. The objective here is to select the inputs’ combinations that lead to coherent states in the internal nodes of the model. To do this, we filter the model’s steady states and select the ones that can reproduce what is known in the literature or observed in transcriptomic datasets. First, we identify the differentially expressed biomolecules present in the model using both literature mining and differential expression analysis (DEA). Then, we discretise the expressions of the identified biomolecules. We convert them to a binary vector of experimentally observed Boolean values. After that, we compare the filtered steady states to this binary vector using a similarity score. We select the states with the highest score and calculate their average vector. The resulting vector represents the calibrated state of the model.

#### 6.1. Identification of differentially expressed biomolecules using literature search and gene expression data analysis

We use low- and high-throughput experimental data to identify the differentially expressed biomolecules in the model under study. First, we thoroughly review the literature regarding each node in the model. We extract information about the change in its expression level between two biological conditions. These conditions are defined based on the system under study. Depending on literature availability, these differential expressions can be at the mRNA and/or protein levels. When it is relevant to the model, we curate the retrieved information to keep it disease- and/or cell-type specific. We integrate transcriptomic dataset(s) as well. We select the dataset(s) according to the biological question we would like the model to address and perform differential expression analysis (DEA) on the selected one(s). The final list of differentially expressed molecules in the model combines literature search and DEA outcomes.

To calibrate the RA M1 and RA M2 macrophage models, we use the GSE97779 dataset, a publicly available microarray dataset from the GEO database^70^. The dataset contains nine RA synovial macrophage samples from nine patients and five peripheral blood monocyte-derived macrophage samples from five healthy donors. We normalised gene expression using quantile normalisation and the preprocessCore package ^71^. We performed DEA using the Limma package ^72^ to identify the DEGs between RA and healthy samples. We filtered the DEGs using an adjusted p-value threshold equal to 0.05.

#### 6.2. Computation of similarity scores and data discretisation

We discretise the data to compare the expressions of the identified differentially expressed components with the model’s steady states. Overexpressed biomolecules in the condition under study are associated with the value one, while under-expressed molecules are associated with the value zero. Since we use Boolean formalism, where each biomolecule can only have two possible states, biomolecules that are not differentially expressed are not considered. The resulting discretised vector of experimentally observed expressions is then used to calculate similarity scores with each steady state to describe the ability of these filtered steady states to reproduce the experimentally observed values. To do so, we calculate a similarity score S (1).

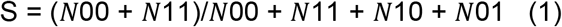

Where,

N00 = number of nodes with a state of zero in both the steady state and the discretised vector of experimentally observed expressions

N01 and N10 = number of nodes with different states in the steady state and the discretised vector of experimentally observed expressions

N11 = number of nodes with a state of one in both the steady state and the discretised vector of experimentally observed expressions

#### 6.3. Selection of the steady states with the highest similarity score

We select the steady states with the highest similarity score. Then, we compute the mean value of each node over these stable states to determine the nodes that are fixed at either zero or one and those that can be found in both states. The resulting average vector represents the calibrated model’s state.

## Data availability

This published article and its supplementary information files include all data generated or analysed during this study.

## Code availability

Scripts and files generated are available at: https://gitlab.com/genhotel/Large_scale_computational_modelling_of_the_M1_and_M2_synovial_macrophages_in_Rheumatoid_Arthritis

Models are available through a review account in BioModels https://www.ebi.ac.uk/biomodels/login/auth:

1. Log in with username reviewerForMODEL2307180001 and password 8X9PYO to have access to the RA M1 macrophage model
2. Access the model at this link https://www.ebi.ac.uk/biomodels/MODEL2307180001
3. Log in with username reviewerForMODEL2307180002 and password Y3SIBT to have access to the RA M2 macrophage model
4. Access the model at this link https://www.ebi.ac.uk/biomodels/MODEL2307180002

## Acknowledgements

The authors would like to thank Dr Sylvain Soliman for his insight and expertise, which considerably supported the work.

This work has been supported by the public-private partnership grant (CIFRE contract, no 2020/0766). B.A.H. acknowledges support from the Royal Society (grant no. UF130039).

## Author contributions

N.Z. performed computational modeling, analyzed data, drafted the manuscript, and revised the manuscript.

R.A. and B.A.H. developed the Linux version of the BMA tool and revised the manuscript.

F.A. supervised the work, analyzed data, drafted the manuscript, and revised the manuscript.

A.N. conceived the study, supervised the work, analyzed data, drafted the manuscript, and revised the manuscript.

All authors revised and approved the manuscript.

