## Supplementary material for "Large-scale computational modelling of the M1 and M2 synovial macrophages in Rheumatoid Arthritis"

**Supplementary Data**

**Table S-1**. The list of differentially expressed genes present in the RA M1 macrophage model that we identified using literature search and omics data analysis. The first column contains the DEGs HGNC names. The second and fifth columns contain their corresponding Boolean values observed in GSE97779 dataset and literature respectively.

| **DEG** | **Boolean value in GSE97779** | **Adjusted p_value** | **logFC** | **Boolean value based on literature** | **Reference** |
| --- | --- | --- | --- | --- | --- |
| C5A |  |  |  | 1 | 29220376 |
| PRKCD | 1 | 0,0008 | 0,86 |  |  |
| BAD | 0 | 0,02492 | -0,42 |  |  |
| STAT2 |  |  |  | 1 | 27626941 |
| IRF9 | 1 | 0,00987 | 0,65 | 1 | 27626941 |
| INHBA |  |  |  | 1 | 13130463 |
| SMAD4 | 0 | 0,012 | -0,46 |  |  |
| CASP1 | 1 | 0,03031 | 0,71 |  |  |
| SIRT1 | 0 | 0,0284 | -0,61 | 0 | 25799392 |
| CCL21 |  |  |  | 1 | 21225692 |
| PRKCQ | 1 | 0,0037 | 1,53 |  |  |
| INHBB |  |  |  | 1 | 26359667 |
| TRAF6 | 0 | 0,00612 | -0,73 |  |  |
| SMAD7 | 1 | 7,8882E-05 | 2,17 |  |  |
| INPP5A | 1 | 0,0091 | 0,9 |  |  |
| DUSP1 | 1 | 5,59E-07 | 4,17 |  |  |
| PRKG1 | 1 | 0,00311 | 2,17 |  |  |
| ACVR2A | 0 | 0,01746 | -0,79 |  |  |
| ACVR2B | 0 | 0,00045 | -2,85 |  |  |
| BCL2 | 1 | 0,01417 | 1,67 |  |  |
| BCL2L1 |  |  |  | 1 | 28118944 |
| BCL2L11 | 1 | 0,0833 | 1,78 |  |  |
| BCL3 | 1 | 0,04434 | 0,42 |  |  |
| BIRC2 | 1 | 0,00425 | 0,594 |  |  |
| C5AR1 |  |  |  | 1 | 29220376 |
| CASP3 | 0 | 0,00542 | -0,86 |  |  |
| CASP7 | 1 | 0,00655 | 0,96 |  |  |
| CCL2 |  |  |  | 1 | 33330982 |
| CCR2 | 1 | 7,69E-07 | 5,85 |  |  |
| CFLAR | 1 | 0,0029 | 0,8 | 1 | 12228167 |
| CSF2RA | 1 | 0,00287 | 1,11 | 1 | 24936585 |
| CSF2RB | 1 | 0,00099 | 0,93 | 1 | 24936585 |
| CXCL1 |  |  |  | 1 | 7561066 |
| FAS | 1 | 0,0005 | 2,45 |  |  |
| FOS | 1 | 2,99E-08 | 3,93 | 1 | 27626941 |
| GNA13 | 1 | 0,00017 | 1,62 |  |  |
| GNAI3 | 1 | 0,00098 | 0,72 |  |  |
| GNB1 | 1 | 0,00507 | 0,47 |  |  |
| HLA-B | 1 | 0,02221 | 0,42 |  |  |
| HRAS | 1 | 0,00388 | 0,94 |  |  |
| IFNA1 | 1 | 0,00831 | 2,43 |  |  |
| IFNB |  |  |  | 1 | 15878901 |
| IFNGR1 | 1 | 0,01789 | 1,35 | 1 | 25708927 |
| IFNGR2 | 1 | 0,00415 | 0,67 | 1 | 25708927 |
| IL11 |  |  |  | 1 | 29327326 |
| IL12RB1 | 1 | 0,00343 | 1,29 |  |  |
| IL18 |  |  |  | 1 | 10562301 |
| IL18R1 | 1 | 0,00057 | 3,64 |  |  |
| IL1RAP | 1 | 0,00018 | 2,73 |  |  |
| IL23 |  |  |  | 1 | 25799392 |
| IL6 | 1 | 0,02477 | 1,56 |  |  |
| IL6ST | 1 | 0,01567 | 0,56 |  |  |
| IL8 |  |  |  | 1 | 10491366 |
| IRF7 | 1 | 1,89337E-05 | 2,44 | 1 | 22614743 |
| JAK1 | 1 | 0,00433 | 1 |  |  |
| JAK2 | 1 | 0,0011 | 2,3 |  |  |
| LILRB1 | 1 | 0,0015 | 0,96 |  |  |
| MAP2K1 | 1 | 0,0001 | 1,12 |  |  |
| MAP2K2 | 1 | 0,00845 | 0,64 |  |  |
| MAP2K3 | 1 | 0,00258 | 1,03 |  |  |
| MAP2K6 | 1 | 0,02686 | 1,78 |  |  |
| MAPK14 | 0 | 0,0243 | -0,61 |  |  |
| MAPK3 |  |  |  | 1 | 17907188 |
| MAPK8 | 1 | 0,00374 | 0,71 |  |  |
| MAPKAPK2 | 1 | 0,00399 | 0,78 |  |  |
| MCL1 | 1 | 0,00118 | 1,39 | 1 | 17009247 |
| MDM2 | 0 | 0,01706 | -1,06 |  |  |
| MYC | 1 | 0,00027 | 1,66 |  |  |
| NFAT5 | 1 | 0,01494 | 0,97 |  |  |
| NFKB1 | 1 | 0,00235 | 0,89 | 1 | 8630106 |
| NFKBIA | 1 | 0,00021 | 0,98 |  |  |
| NFKBIE | 1 | 0,01037 | 0,79 |  |  |
| NLRP3 | 1 | 8,50E-06 | 4,44 |  |  |
| OPN3 | 0 | 0,00517 | -0,87 |  |  |
| PPIA | 1 | 0,01626 | 1,05 |  |  |
| PTK2 | 1 | 0,00073 | 1,87 |  |  |
| PTPN6 | 1 | 2,86974E-05 | 1 |  |  |
| RAC1 | 1 | 0,03366 | 0,44 |  |  |
| RAF1 | 1 | 0,01113 | 1 |  |  |
| RBPJ | 1 | 3,82155E-05 | 2,5 |  |  |
| RELA | 1 | 0,01152 | 0,66 |  |  |
| SMAD2 | 0 | 0,01636 | -0,45 |  |  |
| SOS1 | 1 | 0,01592 | 1,32 |  |  |
| STAT1 | 1 | 0,00057 | 2,22 | 1 | 22614743 |
| STAT3 | 1 | 0,00043 | 0,89 |  |  |
| STAT4 |  |  |  | 1 | 10779770 |
| STAT5B | 1 | 0,00037 | 2,34 |  |  |
| SYK | 0 | 0,03668 | -0,38 |  |  |
| TLR1 | 1 | 1,21E-06 | 1,65 |  |  |
| TLR2 | 1 | 8,13E-07 | 2,2 | 1 | 15146415 |
| TLR4 | 0 | 0,01061 | -0,67 |  |  |
| TLR5 | 0 | 0,00032 | -1,25 |  |  |
| TLR8 | 1 | 0,04753 | 0,68 |  |  |
| TLR9 |  |  |  | 1 | 26759164 |
| TNF | 1 | 7,84837E-05 | 2,5 | 1 | 2109776 |
| TNFRSF11 | 1 | 0,01287 | 1,79 |  |  |
| TNFRSF1A |  |  |  | 1 | 9189061 |
| TRADD | 1 | 0,03339 | 0,47 |  |  |
| TRAF3 | 0 | 0,00022 | -1,14 |  |  |
| TRAM1 | 0 | 0,01224 | -0,4 |  |  |
| XIAP | 1 | 0,00215 | 1,12 | 1 | 19171073 |
| PPP4C | 1 | 0,00131 | 0,68 |  |  |
| biglycan_simple_molecule |  |  |  | 1 | 19772831 |
| DNA_simple_molecule |  |  |  | 1 | 19772831 |
| dsRNA_simple_molecule |  |  |  | 1 | 19772831 |

**Table S-2.** List of nodes upstream the phenotypes of interest in the RA M1 macrophage model associated with their mean values over the fixpoints having the highest similarity score.

| **Nodes** | **Mean values** |
| --- | --- |
| CSF2RA_CSF2RB_complex | 1 |
| RELA_NFKB1_NFKBIE_complex | 1 |
| TLR5 | 0 |
| IRF9 | 1 |
| STAT2 | 1 |
| IKK1_phosphorylated | 0,5 |
| col4a4 | 0,66666667 |
| PTPN6 | 1 |
| PRKG1 | 1 |
| ACVR2A_ACVR2B_complex | 0 |
| ACVR2A_ACVR2B_INHBA_complex | 0 |
| ACVR2A_ACVR2B_INHBB_complex | 0 |
| AP_1 | 1 |
| AP_1_phosphorylated | 1 |
| apoptosis_M1_macrophage_phenotype | 0 |
| ASC | 1 |
| ASK1 | 1 |
| BAD | 0 |
| BCL2_M1_macrophage__Mitochondria_membrane | 1 |
| BCL2_M1_macrophage__Mitochondria_membrane_active | 0 |
| Bcl2_rna | 1 |
| BCL2L1_M1_macrophage__Mitochondria | 1 |
| BCL2L1_M1_macrophage__Mitochondria_active | 0 |
| BCL3_rna | 1 |
| biglycan_simple_molecule | 1 |
| Bim | 1 |
| c_FOS_M1_macrophage___Cytoplasm | 1 |
| c_FOS_M1_macrophage___Cytoplasm_active | 1 |
| c_FOS_M1_macrophage___nucleus | 1 |
| c_JUN | 1 |
| c_JUN_phosphorylated_M1_macrophage___Cytoplasm | 1 |
| c_JUN_phosphorylated_M1_macrophage___nucleus | 1 |
| c_Myc_rna | 1 |
| c5a | 1 |
| C5a_C5aR1_complex | 1 |
| Casp1 | 1 |
| CASP3 | 0 |
| CASP7 | 0 |
| CASP8 | 1 |
| CCL2_CCR2_complex | 1 |
| CCL2_M1_macrophage___Extracellular_Space | 1 |
| CCL2_M1_macrophage___Secreted_components | 1 |
| CCl21 | 1 |
| CCL21_CCR7_complex | 1 |
| CD40LG_ITGB1_ITGA1_complex | 0,5 |
| cFLIP | 1 |
| cIAP1 | 1 |
| col4a5 | 0,66666667 |
| CRKL_phosphorylated | 1 |
| CSF2_M1_macrophage___Extracellular_Space | 1 |
| CSF2_M1_macrophage___Secreted_components | 1 |
| CSF2RA_CSF2RB_CSF2_complex | 1 |
| CXCL1_CXCR1_complex | 1 |
| CXCL1_M1_macrophage___Extracellular_Space | 1 |
| CXCL1_M1_macrophage___Secreted_components | 1 |
| CXCR1 | 1 |
| CXCR1_IL8_complex | 1 |
| CypA | 1 |
| DAXX | 1 |
| DNA_simple_molecule | 1 |
| dsRNA_simple_molecule | 1 |
| DUSP1 | 1 |
| ECSIT | 0,5 |
| ERK1_phosphorylated_M1_macrophage___Cytoplasm | 1 |
| ERK1_phosphorylated_M1_macrophage___nucleus | 1 |
| FADD | 1 |
| FAS | 1 |
| FASL_FAS_complex | 1 |
| FASL_M1_macrophage___Extracellular_Space | 1 |
| FASL_M1_macrophage___Secreted_components | 1 |
| FOXO_M1_macrophage___Cytoplasm | 1 |
| FOXO_M1_macrophage___nucleus | 1 |
| gal | 1 |
| GAL_GALR2_complex | 1 |
| gamma_secretase_complex_complex | 1 |
| GNA12_GNA13_complex | 1 |
| GNAI3 | 1 |
| GNB_GNG_GNAI3_complex | 1 |
| HLA_B_LILRB1_complex | 1 |
| HRAS | 1 |
| IFNa_M1_macrophage___Extracellular_Space | 1 |
| IFNa_M1_macrophage___Secreted_components | 1 |
| IFNAR1_IFNAR2_complex | 1 |
| IFNAR1_IFNAR2_IFNa_complex | 1 |
| IFNAR1_IFNAR2_IFNb_complex | 1 |
| IFNb_M1_macrophage___Extracellular_Space | 1 |
| IFNb_M1_macrophage___Secreted_components | 1 |
| IFNE | 0,5 |
| IFNE_IFNAR1_IFNAR2_complex | 0,5 |
| IFNg_M1_macrophage___Extracellular_Space | 1 |
| IFNg_M1_macrophage___Secreted_components | 1 |
| IFNGR1_IFNGR2_complex | 1 |
| IFNGR1_IFNGR2_IFNg_complex | 1 |
| IKK_complex | 1 |
| IKK1_IKK2_complex | 0,5 |
| IKK2_phosphorylated | 1 |
| IKKE_TBK1_complex | 0,5 |
| IKKE_TBK1_TRAF3_complex | 0 |
| IL1_IL1R_complex | 1 |
| IL11_IL11Ra_IL6ST_complex | 1 |
| IL12_M1_macrophage___Extracellular_Space | 1 |
| IL12_M1_macrophage___Secreted_components | 1 |
| IL12RB_complex | 1 |
| IL12RB_IL12_complex | 1 |
| IL18_IL18R1_complex | 1 |
| IL18_M1_macrophage___Cytoplasm | 1 |
| IL18_M1_macrophage___Extracellular_Space | 1 |
| IL18_M1_macrophage___Secreted_components | 1 |
| IL1B_M1_macrophage___Cytoplasm | 1 |
| IL1B_M1_macrophage___Extracellular_Space | 1 |
| IL1B_M1_macrophage___Secreted_components | 1 |
| AKT1 | 0 |
| IL1R_complex | 1 |
| IL23_M1_macrophage___Extracellular_Space | 1 |
| IL23_M1_macrophage___Secreted_components | 1 |
| IL23R_IL12RB1_complex | 1 |
| IL23R_IL12RB1_IL23_complex | 1 |
| IL6_IL6R_IL6ST_complex | 1 |
| IL6_M1_macrophage___Extracellular_Space | 1 |
| IL6_M1_macrophage___Secreted_components | 1 |
| IL8_M1_macrophage___Extracellular_Space | 1 |
| IL8_M1_macrophage___Secreted_components | 1 |
| INHBA | 1 |
| INHBB | 1 |
| INPP5A | 1 |
| IRAK1 | 1 |
| IRAK1_IRAK4_complex | 1 |
| IRAK4_phosphorylated | 1 |
| IRF3_phosphorylated | 1 |
| IRF7 | 1 |
| ITGB1_ITGA1_col4a_complex | 1 |
| ITGB1_ITGA1_complex | 1 |
| JAK1 | 1 |
| JAK1_JAK2_complex | 1 |
| JAK1_TYK2_complex | 1 |
| JAK2 | 1 |
| JAK2_TYK2_complex | 1 |
| JNK1_phosphorylated_M1_macrophage___Cytoplasm | 1 |
| JNK1_phosphorylated_M1_macrophage___nucleus | 1 |
| Mcl1_rna | 1 |
| MDM2_phosphorylated | 0 |
| MEK1_phosphorylated | 1 |
| MEK2_phosphorylated | 1 |
| MEKK1 | 0,5 |
| MK2_phosphorylated | 1 |
| MKK3_phosphorylated | 1 |
| MKK4_phosphorylated | 1 |
| MKK6_phosphorylated | 1 |
| MKK7_phosphorylated | 1 |
| MYD88 | 1 |
| MYD88_TIRAP_TOLLIP_complex | 1 |
| NCID | 1 |
| NFAT5 | 1 |
| NFKB1_TPL2_complex | 1 |
| NFKBIA_RELA_NFKB1_complex | 1 |
| NICD | 1 |
| NICD_CSL_SKIP_MAML1_ep300_complex | 1 |
| NIK | 1 |
| NLRP3 | 1 |
| NLRP3_INFLAMMASOME_complex | 1 |
| notch1_JAG1_complex | 1 |
| OPN | 0 |
| osteoclastogenesis_M1_macrophage_phenotype | 1 |
| p15_rna | 0 |
| p21_rna | 1 |
| p300_SP1_complex | 0 |
| p38_MAP_KINASE_phosphorylated_M1_macrophage___Cytoplasm | 0 |
| p38_MAP_KINASE_phosphorylated_M1_macrophage___nucleus | 0 |
| p53_phosphorylated | 1 |
| PI3K | 0 |
| PIK3AP1_phosphorylated | 0 |
| PP4 | 1 |
| Prkcd | 1 |
| PRKCQ | 1 |
| proliferation_survival_M1_macrophage_phenotype | 1 |
| PTK2 | 1 |
| Rac1 | 1 |
| RAF1 | 1 |
| RELA_NFKB1_complex_M1_macrophage___Cytoplasm | 1 |
| RELA_NFKB1_complex_M1_macrophage___nucleus | 1 |
| RHOA | 0 |
| SHP2_GRB2_complex | 1 |
| Sirt1 | 0 |
| SMAD2_phosphorylated | 0 |
| SMAD2_SARA_complex | 0 |
| SMAD2_SMAD4_complex | 0 |
| SMAD4 | 0 |
| SMAD7 | 1 |
| SOS1 | 1 |
| Src | 1 |
| STAT1 | 1 |
| STAT1_STAT1_complex | 1 |
| STAT1_STAT2_IRF9_complex | 1 |
| STAT3 | 1 |
| STAT3_STAT3_complex | 1 |
| STAT4 | 1 |
| STAT4_STAT4_complex | 1 |
| STAT5_CRKL_complex | 1 |
| STAT5_phosphorylated | 1 |
| SYK | 0 |
| TAB1 | 1 |
| TAB2_phosphorylated | 1 |
| TAK1 | 1 |
| TLR1_TLR2_biglycan_complex | 1 |
| TLR1_TLR2_complex | 1 |
| TLR2_TLR6_biglycan_complex | 1 |
| TLR2_TLR6_complex | 1 |
| TLR3_dsRNA_complex | 1 |
| TLR4_Md2_CD14_fibrinogen_complex | 0 |
| TLR7_TLR8_ssRNA_complex | 1 |
| TLR9_DNA_complex | 1 |
| TNF_M1_macrophage___Extracellular_Space | 1 |
| TNF_M1_macrophage___Secreted_components | 1 |
| TNF_TNFRSF1A_complex | 1 |
| TNFA_rna | 1 |
| TNFSF11 | 1 |
| TNFSF11_TNFRSF11_complex | 1 |
| TPL2 | 1 |
| TRADD | 1 |
| TRADD_TRAF2_RIP1_complex | 1 |
| TRAF2_RIP1_TRADD_TAK1_TAB1_TAB2_complex | 1 |
| TRAF2_TRAF6_complex | 1 |
| TRAF3 | 0 |
| TRAF6 | 0 |
| TRAF6_ECSIT_MEKK1_TAB1_TAB2_TAK1_complex | 0 |
| TRAF6_TAB1_TAB2_TAK1_complex | 0 |
| TRAF6_ubiquitinated | 0 |
| TRAM1 | 0 |
| TRAM1_TRIF_complex | 0 |
| TRIF | 1 |
| TSG6 | 1 |
| UEV1A_UBC13_complex | 0,5 |
| XIAP | 1 |

**Table S-3**. The list of differentially expressed genes present in the RA M2 macrophage model that we identified using literature search and omics data analysis. The first column contains the DEGs HGNC names. The second and fifth columns contain their corresponding Boolean values observed in GSE97779 dataset and literature respectively.

| **DEG** | **Boolean value in GSE97779** | **Adjusted p_value** | **logFC** | **Boolean value based on literature** | **Reference** |
| --- | --- | --- | --- | --- | --- |
| VEGFB | 0 | 9,64E-06 | -1,46 |  |  |
| PRLR | 0 | 0,01306 | -1,45 |  |  |
| MDM2 | 0 | 0,01706 | -1,06 |  |  |
| CASP3 | 0 | 0,00542 | -0,86 |  |  |
| TRAF6 | 0 | 0,00612 | -0,73 |  |  |
| CREB1 | 0 | 0,03541 | -0,66 |  |  |
| NRP2 | 0 | 0,03197 | -0,65 |  |  |
| MAPK14 | 0 | 0,0243 | -0,61 |  |  |
| SIRT1 | 0 | 0,0284 | -0,61 | 0 | 25799392 |
| SYK | 0 | 0,015846 | -0,53 |  |  |
| SMAD4 | 0 | 0,012 | -0,46 |  |  |
| SMAD2 | 0 | 0,01636 | -0,45 |  |  |
| BAD | 0 | 0,02492 | -0,42 |  |  |
| BCL3 | 1 | 0,04434 | 0,42 |  |  |
| HLA-B | 1 | 0,02221 | 0,42 |  |  |
| CEBPB | 1 | 0,00957 | 0,43 |  |  |
| SHC1 | 1 | 0,03891 | 0,44 |  |  |
| STAT6 | 1 | 0,02689 | 0,44 |  |  |
| BAX | 1 | 0,02724 | 0,45 | 1 | 12634940 |
| GNB1 | 1 | 0,00507 | 0,47 |  |  |
| IL6ST | 1 | 0,01567 | 0,56 |  |  |
| CYLD | 1 | 0,01375 | 0,61 |  |  |
| FCGR2A | 1 | 0,02327 | 0,63 | 1 | 17521421 |
| MAP2K2 | 1 | 0,00845 | 0,64 |  |  |
| RELA | 1 | 0,01152 | 0,66 |  |  |
| CASP1 | 1 | 0,03031 | 0,71 |  |  |
| GNAI3 | 1 | 0,00098 | 0,72 |  |  |
| TGFB1 | 1 | 0,00044 | 0,78 |  |  |
| NFKBIE | 1 | 0,01037 | 0,79 |  |  |
| CFLAR | 1 | 0,0029 | 0,8 | 1 | 12228167 |
| RXRA | 1 | 0,0023 | 0,84 |  |  |
| IL4R | 1 | 0,00047 | 0,86 | 1 | 7492352 |
| PRKCD | 1 | 0,0008 | 0,86 |  |  |
| NFKB1 | 1 | 0,00235 | 0,89 | 1 | 8630106 |
| STAT3 | 1 | 0,00043 | 0,89 |  |  |
| HRAS | 1 | 0,00388 | 0,94 |  |  |
| CASP7 | 1 | 0,00655 | 0,96 |  |  |
| LILRB1 | 1 | 0,0015 | 0,96 |  |  |
| NFAT5 | 1 | 0,01494 | 0,97 |  |  |
| NFKBIA | 1 | 0,00021 | 0,98 |  |  |
| JAK1 | 1 | 0,00433 | 1 |  |  |
| PTPN6 | 1 | 2,86974E-05 | 1 |  |  |
| RAF1 | 1 | 0,01113 | 1 |  |  |
| SH2D1A | 1 | 0,04778 | 1,01 |  |  |
| IL17RA | 1 | 0,0044 | 1,02 | 1 | 19265168 |
| MAP2K3 | 1 | 0,00258 | 1,03 |  |  |
| EFNB1 | 1 | 0,0094 | 1,09 |  |  |
| MAP2K1 | 1 | 0,0001 | 1,12 |  |  |
| PLCG2 | 1 | 0,00027 | 1,12 |  |  |
| XIAP | 1 | 0,00215 | 1,12 | 1 | 19171073 |
| HCK | 1 | 5,23869E-05 | 1,13 | 1 | 17963503 |
| IL12RB1 | 1 | 0,00343 | 1,29 |  |  |
| SOS1 | 1 | 0,01592 | 1,32 |  |  |
| HBEGF | 1 | 0,04278 | 1,34 | 1 | 31068444 |
| MCL1 | 1 | 0,00118 | 1,39 | 1 | 17009247 |
| LIFR | 1 | 0,01414 | 1,53 |  |  |
| PRKCQ | 1 | 0,0037 | 1,53 |  |  |
| MYC | 1 | 0,00027 | 1,66 |  |  |
| BCL2 | 1 | 0,01417 | 1,67 |  |  |
| BCL2L11 | 1 | 0,0833 | 1,78 |  |  |
| MAP2K6 | 1 | 0,02686 | 1,78 |  |  |
| PTK2 | 1 | 0,00073 | 1,87 |  |  |
| VEGFA | 1 | 0,00429 | 1,9 |  |  |
| SMAD7 | 1 | 7,8882E-05 | 2,17 |  |  |
| JAK2 | 1 | 0,0011 | 2,3 |  |  |
| FAS | 1 | 0,0005 | 2,45 |  |  |
| FCGR3A | 1 | 2,49141E-05 | 2,77 | 1 | 22235253 |
| FCGR1A | 1 | 0,00011 | 3,32 | 1 | 17521421 |
| DUSP1 | 1 | 5,59417E-07 | 4,17 |  |  |
| NLRP3 | 1 | 8,50103E-06 | 4,44 |  |  |
| KLF4 | 1 | 1,77E-08 | 4,66 | 1 | 29997611 |
| BCL2L1 |  |  |  | 1 | 28118944 |
| C5A |  |  |  | 1 | 29220376 |
| C5AR1 |  |  |  | 1 | 29220376 |
| CCL21 |  |  |  | 1 | 21225692 |
| CSF1 |  |  |  | 1 | 27036883 |
| CSF1R |  |  |  | 1 | 27036883 |
| IL10 |  |  |  | 1 | 20001767 |
| IL11 |  |  |  | 1 | 29327326 |
| IL17F |  |  |  | 1 | 19265168 |
| IL17RC |  |  |  | 1 | 19265168 |
| IL23 |  |  |  | 1 | 25799392 |
| IL34 |  |  |  | 1 | 22039170 |
| IL4 |  |  |  | 1 | 7492352 |
| IL8 |  |  |  | 1 | 10491366 |
| MAPK3 |  |  |  | 1 | 17907188 |
| TGFBR1 |  |  |  | 1 | 7614737 |
| TGFBR2 |  |  |  | 1 | 7614737 |

**Table S-4.** List of nodes upstream the phenotypes of interest in the RA M2 macrophage model associated with their mean values over the fixpoints having the highest similarity score.

| **Nodes** | **Mean values** |
| --- | --- |
| AKT1 | 0 |
| AKT1_phosphorylated | 0 |
| apoptosis_M2_macrophage_phenotype | 1 |
| ASC | 0,5 |
| ASK1 | 1 |
| BAD | 0 |
| BAX | 1 |
| BCL2_M2_macrophage_mitochondrion_membrane | 1 |
| BCL2_M2_macrophage_mitochondrion_membrane_active | 0 |
| Bcl2_rna | 1 |
| BCL2L1 | 0 |
| BCL3_rna | 1 |
| Bim | 1 |
| C_EBPb_phosphorylated | 1 |
| c_Myc_rna | 1 |
| c5a | 1 |
| C5a_C5aR1_complex | 1 |
| Casp1 | 1 |
| CASP3 | 1 |
| CASP7 | 1 |
| CASP8 | 1 |
| CASP9 | 0 |
| CCl21 | 1 |
| CCL21_CCR7_complex | 1 |
| CD32a | 1 |
| CD40LG_ITGB1_ITGA1_complex | 0,5 |
| cFLIP | 1 |
| cMyc | 1 |
| cMyc_phosphorylated | 1 |
| col4a4 | 0,66666667 |
| col4a5 | 0,66666667 |
| CREB1_phosphorylated | 0 |
| CSF1_M2_macrophage__extracellular_space | 1 |
| CSF1_M2_macrophage__secreted_components | 1 |
| CSF1R | 1 |
| CSF1R_CSF1_complex | 1 |
| CSFR1R_IL34_complex | 1 |
| CXCR1_IL8_complex | 1 |
| CYLD | 1 |
| DAG_simple_molecule | 1 |
| DAXX | 1 |
| DUSP1 | 1 |
| EFNB1_EPHB1_complex | 1 |
| ERK1_phosphorylated_M2_macrophage__cytoplasm | 1 |
| ERK1_phosphorylated_M2_macrophage_nucleus | 1 |
| FADD | 1 |
| FAS | 1 |
| FASL_FAS_complex | 1 |
| FASL_M2_macrophage__extracellular_space | 1 |
| FASL_M2_macrophage__secreted_components | 1 |
| FOXO1 | 1 |
| GAB2_phosphorylated | 1 |
| GAS6 | 0,5 |
| GAS6_MERTK_complex | 0,5 |
| GNAI3 | 1 |
| GNB_GNG_complex | 1 |
| GNB_GNG_GNAI3_complex | 1 |
| Grb2 | 1 |
| GSK3B | 1 |
| HBGEF | 1 |
| HCK | 1 |
| HLA_B_LILRB1_complex | 1 |
| HRAS | 1 |
| IKK_complex | 1 |
| IKK1_IKK2_complex | 0,5 |
| IKK1_phosphorylated | 0,5 |
| IKK2_phosphorylated | 1 |
| IL10_M2_macrophage__extracellular_space | 1 |
| IL10_M2_macrophage__secreted_components | 1 |
| IL10R1_IL10R2_complex | 1 |
| IL10R1_IL10R2_IL10_complex | 1 |
| IL11Ra_IL6ST_IL11_complex | 1 |
| IL17Ra_IL17Rc_IL17F_complex | 1 |
| IL23_M2_macrophage__extracellular_space | 1 |
| IL23_M2_macrophage__secreted_components | 1 |
| IL23R_IL12RB1_complex | 1 |
| IL23R_IL12RB1_IL23_complex | 1 |
| IL34 | 1 |
| IL4 | 1 |
| IL4_IL4Ra_complex | 1 |
| IL4R | 1 |
| IL8_M2_macrophage__extracellular_space | 1 |
| IL8_M2_macrophage__secreted_components | 1 |
| immune_complex_CD16a_complex | 1 |
| immune_complex_CD32a_complex | 1 |
| immune_complex_CD32b_complex | 1 |
| immune_complex_CD64_complex | 1 |
| immune_complex_complex | 1 |
| ITGB1_ITGA1_col4a_complex | 1 |
| ITGB1_ITGA1_complex | 1 |
| JAK1 | 1 |
| JAK1_TYK2_complex | 1 |
| JAK2 | 1 |
| klf4 | 1 |
| LIFR_IL6ST_CTF1_complex | 1 |
| mcl1 | 0 |
| Mcl1_rna | 0 |
| MDM2_phosphorylated | 0 |
| MEK1_phosphorylated | 1 |
| MEK2_phosphorylated | 1 |
| MKK3_phosphorylated | 1 |
| MKK6_phosphorylated | 1 |
| MSK1_phosphorylated | 0 |
| NFAT5_phosphorylated | 1 |
| NFKB1_TPL2_complex | 1 |
| NFKBIA_RELA_NFKB1_complex | 1 |
| NLRP3 | 1 |
| NLRP3_INFLAMMASOME_complex | 0,5 |
| p15_rna | 1 |
| p38_MAP_KINASE_phosphorylated_M2_macrophage__cytoplasm | 0 |
| p38_MAP_KINASE_phosphorylated_M2_macrophage_nucleus | 0 |
| p53_phosphorylated_M2_macrophage__cytoplasm | 1 |
| p53_phosphorylated_M2_macrophage_nucleus | 1 |
| PI3K | 0 |
| PIK3AP1_phosphorylated | 0 |
| PIP2_simple_molecule | 1 |
| PLCG2 | 1 |
| PRKCD | 1 |
| Prkcd | 1 |
| PRKCQ | 1 |
| PRL | 0 |
| PRL_PRLR_complex | 0 |
| proliferation_survival_M2_macrophage_phenotype | 0 |
| PTK2 | 1 |
| PTPN6 | 1 |
| RAF1 | 1 |
| RBL1_E2F4_DP1_complex | 0 |
| RELA_NFKB1_complex_M2_macrophage__cytoplasm | 1 |
| RELA_NFKB1_complex_M2_macrophage_nucleus | 1 |
| RELA_NFKB1_NFKBIE_complex | 1 |
| RXRa_NUR77_complex | 1 |
| SH2D1A | 1 |
| Shc_phosphorylated | 1 |
| SHIP1 | 1 |
| Sirt1 | 0 |
| SMAD2_phosphorylated | 0 |
| SMAD2_SARA_complex | 0 |
| SMAD2_SMAD4_complex | 0 |
| SMAD4 | 0 |
| SMAD7 | 1 |
| SOS1 | 1 |
| Src | 1 |
| STAT3 | 1 |
| STAT3_STAT3_complex | 1 |
| STAT6 | 1 |
| STAT6_STAT6_complex | 1 |
| SYK | 0 |
| Syk_phosphorylated | 1 |
| TAK1_phosphorylated | 1 |
| TGFB1_M2_macrophage__extracellular_space | 1 |
| TGFB1_M2_macrophage__secreted_components | 1 |
| TGFBR1_TGFBR2_complex | 1 |
| TGFBR1_TGFBR2_TGFB1_complex | 1 |
| TPL2 | 1 |
| TRAF3IP2_phosphorylated | 1 |
| TRAF6_ubiquitinated | 0 |
| VEGFa_M2_macrophage__extracellular_space | 1 |
| VEGFa_M2_macrophage__secreted_components | 1 |
| Vegfa_rna | 1 |
| Vegfb_M2_macrophage__extracellular_space | 0 |
| Vegfb_M2_macrophage__secreted_components | 0 |
| Vegfc_M2_macrophage__extracellular_space | 0 |
| Vegfc_M2_macrophage__secreted_components | 0 |
| vegfc_vegfr3_complex | 0 |
| VegfR1 | 0,5 |
| VegfR1_Vegfa_complex | 0,5 |
| VegfR1_vegfb_complex | 0 |
| VegfR2_vegfc_nrp2_complex | 0 |
| XIAP | 1 |

**Table S-5.** Therapeutic drug targets in the RA M1 macrophage model

**Therapeutic target**

SRC

SIRT1

MAPK14

AKT1

PTPN6

TLR9

TLR8

TLR7

MCL1

MDM2

JAK2

JAK1

CSF2

IL23R

SYK

RAF1

CCR2

BCL2L1

CASP3

CASP8

FAS

ACVR2B

IRAK4

BCL2

C5AR1

TBK1

GALR2

IRAK1

TYK2

CASP1

CASP7

NLRP3

PRKCD

XIAP

PTK2

PPIA

TLR2

TLR4

PRKCQ

IL6R

IL6ST

TNF

DUSP1

JUN

NFKB1

RELA

CXCR1

MAPK3

ITGB1

MYC

STAT3

RAC1

IL1B

CCL2

RHOA

STAT1

IFNG

CD40LG

TNFSF11

IL6

INHBA

INHBB

NOTCH1

CSF2RB

IL12A

ITGA1

NFKBIA

IFNAR2

SMAD7

ACVR2A

IL18

**Table S-6.** Therapeutic drug targets in the RA M2 macrophage model

**Therapeutic Target**

SRC

GSK3B

SIRT1

MAPK14

AKT1

PTPN6

HCK

MCL1

MDM2

JAK2

CSF1R

JAK1

VEGFA

IL23R

IL4R

SYK

RAF1

TGFBR1

BCL2L1

CASP3

CASP8

FAS

BCL2

C5AR1

TYK2

RXRA

MERTK

TGFBR2

CASP1

CASP7

NLRP3

PRKCD

XIAP

PTK2

PRKCQ

IL6ST

CASP9

DUSP1

NFKB1

RELA

CXCR1

MAPK3

ITGB1

MYC

STAT3

CD40LG

KLF4

PRLR

CSF1

TGFB1

LIFR

IL4

ITGA1

GAS6

VEGFC

NFKBIA

BAX

SMAD7

IL17RA

IL17RC
